# *T*_50_ as a proxy of PS II heat tolerance: time dependence, differences in calculation methods and implications for its suitability to assess the heat tolerance of tree species

**DOI:** 10.64898/2026.07.24.740464

**Authors:** Markus Hauck, Choimaa Dulamsuren

## Abstract

- The heat tolerance of tree species has recently received increased attention, as it is critical for the response of forest ecosystems to climate change. Most studies rely on laboratory experiments with detached leaves, where the thermostability of photosystem II (PS II) is analyzed using chlorophyll fluorescence analysis. The temperature at which the maximum quantum yield (*F*_v_/*F*_m_) of PS II is reduced by 50% (*T*_50_) is often used as key measure of heat tolerance despite of weak mechanistic corroboration and inconsistent definitions. We propose a new metric consisting of a critical temperature (*T*_IP_) and a corresponding fluorescence value (*F*_IP_).
- We analyzed the effect of incubation time and temperature on different versions of *T*_50_.
- *T*_50_ was strongly dependent on the duration of heat exposure, decreasing exponentially with incubation time. *T*_50_ and *T*_IP_ values derived from short-term treatments of different incubation times <1 h are not comparable, but stabilize after longer intervals of heat exposure.
- *T*_50_ of *F*_v_/*F*_m_ specified without considering incubation time is a meaningless metric. Specifications derived from short-term heat treatments should be avoided, because the result is highly influenced by the experimental setup. Different calculation methods for *T*_50_ influence the result and elucidate different aspects of the heat response.

## 1. Introduction

Heat has long been underestimated as a critical factor affecting tree vitality under climate change (Hauck et al. 2025; Posch et al. 2025) compared with the widely acknowledged role of drought (Groover et al. 2025). Reductions in CO_2_ assimilation represent an early, but reversible heat response, which is caused by the inactivation of Rubisco activase even below 40°C (Salvucci & Crafts-Brandner 2004; Haldimann & Feller 2005). Its effect on the tree’s carbon balance is aggravated by increased respiration at high temperatures (Teskey et al. 2015). At somewhat higher temperatures, the light reaction of photosynthesis is impaired, at first reversibly and then irreversibly (Didion-Gency et al. 2025), because the photosystem II (PS II) is markedly heat- sensitive (Allakhverdiev et al. 2008). PS II dysfunction is caused by a cascade of heat-induced ultrastructural and biochemical changes in the chloroplast and can be reliably tracked using chlorophyll fluorescence analysis. Since the repair mechanisms of PS II are even more heat- sensitive than the PS II itself, strong damage of the PS II is assumed to be largely irreversible (Mohanty et al. 2012). Because the light-reaction of photosynthesis is essential, widespread PS II dysfunction should inevitably lead to whole-plant death (Guadagno et al. 2017), at least if it occurs during protracted compound heat and drought periods and non-structural carbohydrate (NSC) reserves are limited (O’Brien et al. 2014; Dulamsuren et al. 2023).

While there is broad agreement about these principles, studies on the PS II heat tolerance of trees differ in the experimental setups and in the interpretation of the results. Most studies on the heat tolerance of trees rely on the exposure of detached leaves or leaf disks to heat in a water bath (Table 1). However, individual researchers followed different strategies with respect to incubation time. The vast majority of studies preferred short-term heat treatments of 15 or 30 min (Table 1) following the argument of Krause et al. (2010) that longer heat exposures of leaves or leaf disks that are encapsulated in a plastic bag and then stored in a water bath could suffer from anaerobiosis. Yet, recent tests of up to 4 h (Hauck et al. 2025) or even 8.5 h (Neuner & Buchner 2023) demonstrated that these concerns were unsubstantiated, which opened the door for experiments with more realistic time spans of heat exposure that come close to the length of the hot noon and afternoon hours during a heat wave. Some recent experiments with tree species from boreal (Dulamsuren et a. 2026), temperate (Hauck et al. 2026), and subtropical (Aitken et al. 2026) forests covered variable incubation periods of up to 2 h or 4 h (Table 1). Kunert et al. (2026b) also employed heat treatments of up to 4 h for Mediterranean tree species, but assumed that the most relevant damage during heat waves is likely to occur during short-term heat maxima. This view agrees with studies in desert plants, where short transient lulls can lead to rapid increases of leaf temperatures within few minutes (Curtis et al. 2014). Curtis et al. (2014) reported short-term temperature peaks above 52°C that are indeed fundamentally critical to a wide range of metabolic processes and to ultrastructural integrity, but these results from an ecosystem with low significance of latent heat are hardly transferable to forest ecosystems with lower leaf temperatures (Leuzinger & Körner 2007; Manzi et al. 2025; Dulamsuren et al. 2026) and higher thermal buffer capacity due to transpiration and the cooling influence of the shaded air masses below the canopy (Adhikari et al. 2025). Stable compound heat and drought periods, like for instance in the summers of 2003, 2018, and 2026 in the temperate forest region of Western and Central Europe, however, demonstrated that hot weather conditions with physiologically critical temperatures can last for days and intensify under the conditions of climate change (Becker et al. 2022).

**Table 1.**
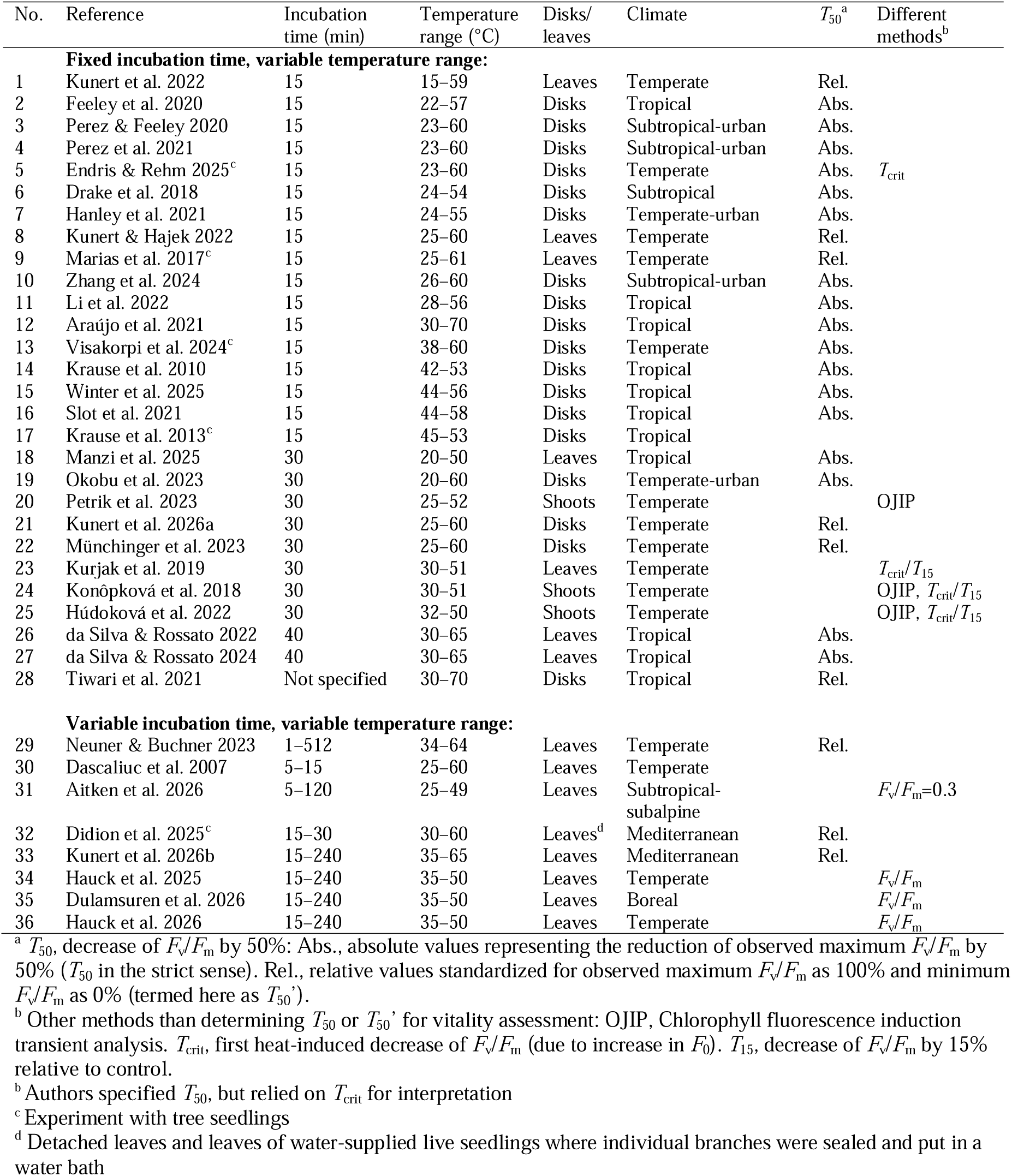
Examples of experimental studies of PS II heat tolerance in tree species after exposure of leaves or leaf disks to different temperatures.

In addition to variations in the experimental methodologies, there are different approaches in the use of chlorophyll fluorescence parameters for the quantification of heat damage. The parameter most widely used in studies of tree species is *T*_50_ (Table 1), which is defined as the temperature at which the maximum quantum yield (*F*_v_/*F*_m_) of PS II is reduced by 50% (Krause et al. 2010; Perez & Feeley 2020). However, since *T*_50_ depends on two parameters, namely temperature and incubation time (Neuner & Buchner 2023), and the duration of heat exposure varies between individual studies (Table 1), its use has been avoided in some recent studies (Hauck et al. 2025, 2026; Aitken et al. 2026; Dulamsuren et al. 2026).

Taking a closer look at the studies using *T*_50_ in the context of PS II heat tolerance, it becomes evident that authors employed different approaches for calculating *T*_50_ (Table 1). Most studies use the maximum *F*_v_/*F*_m_ at the lowest tested temperature as the initial value for the calculation and take the temperature value at which *F*_v_/*F*_m_ is reduced to 50% of this initial value as *T*_50_ (e.g. Krause et al. 2010; Perez & Feeley 2020). If we talk of *T*_50_ in the following, we always mean this definition of *T*_50_, which represents the absolute variant of *T*_50_ (or as one could also say, *T*_50_ in the strict sense). A minority of studies standardizes *F*_v_/*F*_m_ before the calculation of *T*_50_ by setting the maximum *F*_v_/*F*_m_ at low temperature to 100% (or 1) and the lowest *F*_v_/*F*_m_ at high temperature to 0% (or 0) (e.g. Marias et al. 2017; Tiwari et al. 2021; Kunert et al. 2026a, b). We propose to term this relative variant of *T*_50_ as *T*_50_’ and will consistently refer to *T*_50_’ for this relative parameter and to *T*_50_ for the absolute parameter in the rest of this paper.

Both *T*_50_ and *T*_50_’ are valid parameters and the authors of all studies cited in Table 1 provided proper descriptions in their papers of how they determined the temperature corresponding to 50% loss of *F*_v_/*F*_m_. Yet to the best of our knowledge, it has never been explicitly noted in the literature that there are these alternative ways of calculation. Nor have the implications been discussed with respect to differences in the interpretation of *T*_50_ and *T*_50_’ and how these differences may render the use of one of these parameters more recommendable than the other. With the present paper, we wish to fill this gap in addition to the study of the influence of incubation time on *T*_50_ and *T*_50_’.

The function of *F*_v_/*F*_m_ versus temperature at a given incubation time generally follows a sigmoid curve with quite stable *F*_v_/*F*_m_ at low temperatures, a more or less rapid decline in a critical temperature range, and finally *F*_v_/*F*_m_ values of or near zero beyond that temperature threshold (Slot et al. 2021; Posch et al. 2025). If *F*_v_/*F*_m_ values of zero that represent the complete collapse of PS II are reached, depends on the heat tolerance of the studied species and the temperature range investigated. While some tree species reach zero values of *F*_v_/*F*_m_ at temperatures around 45°C, there are other species, where 50°C or even higher temperatures are necessary to produce such effect (Perez & Feeley 2020; da Silva & Rossatto 2022; Visakorpi et al. 2024; Hauck et al. 2025). The initial value of *F*_v_/*F*_m_ is ideally around 0.83 (Murchie & Lawson 2013), but usually also somewhat lower values of ≥0.75 are accepted, because leaves can be exposed to diverse stresses in the field before sampling, but also due to their treatment after sampling, even though experimenters try of course to treat their plant samples as perfectly as possible (Kunert et al. 2026b).

Based on the sigmoid shape of the relationship of *F*_v_/*F*_m_ versus temperature, Didion-Gency et al. (2025) employed a third version of *T*_50_ in the wide sense, which is related to, but not identical with *T*_50_’. While the maximum and minimum values of *F*_v_/*F*_m_ used for the standardization of the chlorophyll fluorescence data equal the upper and lower asymptotes of the sigmoid function in *T*_50_’ (Perez et al. 2021; Kunert et al. 2026b), Didion-Gency et al. (2025) selected a lower reference for maximum *F*_v_/*F*_m_ and a higher reference for minimum *F*_v_/*F*_m_. For the upper limit, the authors chose the critical temperature (*T*_crit_), which is another widely used parameter of PS II heat tolerance and is defined as the temperature, where first heat-induced decreases in *F*_v_/*F*_m_ are detectable, primarily driven by an increase in minimum fluorescence at dark (*F*_0_) due to initial membrane damage, which reduces variable fluorescence (*F*_v_), which is calculated as the difference of maximum fluorescence *F*_m_ and *F*_0_ (Hüve et al. 2011). Often a decrease of in *F*_v_/*F*_m_ by 15% is taken for the definition of *T*_crit_ (Perez & Feeley 2020; Endris & Rehm 2025), while sometimes *T*_crit_ is more qualitatively defined as the temperature where the first drop of *F*_v_/*F*_m_ (corresponding to an increase in *F*_0_) is observed (Krause et al. 2010; Konôpková et al. 2018; Didion-Gency et al. 2025).

For the lower limit, Didion-Gency et al. (2025) defined an *F*_v_/*F*_m_ value above the absolute minimum at the point of the sigmoid curve, where the steep slope inclination transitions into low slope inclination. This procedure is similar to that for determining *T*_50_’, but not identical. Therefore, we term the resulting parameter *T*_50_”. Didion-Gency et al. (2025) called the temperature value that marked their lower temperature limit *T*_max_ and referred for this parameter to Perez et al. (2021). However, although the study of Didion-Gency et al. (2025) represents seminal work on the reversibility of heat damage and on the validity of results obtained with detached leaves versus intact plants, their definition of *T*_max_ is incorrect, because Perez et al. (2021) related *T*_max_ to the dark reaction of photosynthesis and defined *T*_max_ as the high temperature compensation point of CO_2_ gas exchange. Since *T*_max_ as applied by Perez et al. (2021) is well defined and different from the point Didion-Gency et al. (2025) mean, we discourage to use *T*_50_” to avoid ambiguities and also because we do not see an advantage of using *T*_50_” compared to the more widely used *T*_50_’. We will, therefore, not deal with *T*_50_” when comparing parameters in the remaining paper, but would like to emphasize that the use of *T*_50_” did not affect the high value of the paper of Didion-Gency et al. (2025) for progress in PS II heat tolerance research.

Initial *F*_v_/*F*_m_ values around 0.8 in untreated leaves imply that *T*_50_ is always found at *F*_v_/*F*_m_ values near 0.4. *T*_50_ can or cannot be located in the curve section of rapid decline of *F*_v_/*F*_m_ (Krause et al. 2010, 2013; Araujo et al. 2021; da Silva & Rossatto 2022). This illustrates that the selection of *T*_50_ as a key indicator of heat tolerance is arbitrary and not supported by any mechanistic explanation. Even if the exact mechanisms are not known, the curve section with the steepest slope of the regression line for *F*_v_/*F*_m_ versus temperature indicates that the functionality of PS II is rapidly reduced at small increases in temperature in this temperature range and thus suggests that the temperature at which the slope inclination is steepest is critical. There is little evidence whether *F*_v_/*F*_m_ values around 0.4 represent a critical state that is irreversible and would thus be critical for long-term survival. While Slot et al. (2021) and Endris & Rehm (2025) assumed already heat-induced decreases of *F*_v_/*F*_m_ above 15% (i.e. at *T*_crit_) to reflect irreversible damage, there is reason for more careful assessments, because the reversibility of heat-induced PS II damage is poorly studied (Posch et al. 2025). Examining reversibility of *T*_crit_ in three tree species experimentally, Didion-Gency et al. (2025) considered *T*_crit_ to be largely reversible. Assuming that *T*_crit_ represents the first signs of mainly reversible heat damage due to increased membrane fluidity (Hüve et al. 2011; Mohanty et al. 2012), the term “critical temperature” (*T*_crit_) might be somewhat misleading from its literal meaning (e.g. van Tiel et al. 2026) and could lead to overinterpretation, because *T*_crit_ does certainly not represent a lethal temperature (Didion-Gency et al. 2025).

Several studies with short-term exposures to heat allowed the leaf or leaf disk samples to recover 1 day prior to chlorophyll fluorescence analysis (Curtis et al. 2014; Drake et al. 2018; Feeley et al. 2020; Perez & Feeley 2020; Hanley et al. 2021; Perez et al. 2021; Kunert & Hajek 2022; Kunert et al. 2022, 2026a, b; Slot et al. 2021; Visakorpi et al. 2024). However, since all these studies have in common that *F*_v_/*F*_m_ was only determined the day after heat exposure, the results to do not allow for any conclusions regarding potential recovery after the treatment. In the early work of Krause et al. (2010), which is often referred to as a key reference for the methodology in these studies, chlorophyll fluorescence was therefore measured immediately after the heat treatment (following dark adaptation) and 1 day later. Recovery of *F*_v_/*F*_m_ during this day in *Ficus insipida* was minor with mostly overlapping standard deviations suggesting the lack of a significant difference, which was not explicitly tested (Krause et al. 2010). After a 2-min heat treatment in two African tropical tree species, Manzi et al. (2025) observed an increase in *T*_50_ by 2.7 K (*Prunus africana*) and 3.8 K (*Maesa lanceolata*) after 1 day of recovery. In five neotropical tree species, no change in *T*_50_’ was recorded after 1 day of recovery (Tiwari et al. 2021).

The ability of plants to recover from temperature stress is influenced by light conditions (Kreslavski et al. 2008), which varied in the experiments with trees between low light (e.g. Dascaliuc et al. 2007; Feeley et al. 2020; Münchinger et al. 2023) and complete dark (Drake et al. 2018; Hanley et al. 2021; Manzi et al. 2025). Chances for recovery at dark (and under strong illumination) are generally limited (Havaux et al. 1991; Kreslavski et al. 2008). In addition, it might be questioned whether “recovery” as implemented in the experiments always had the potential to have a regenerative effect on the plants. In most cases, leaf disks (e.g. Drake et al. 2018; Feeley et al. 2020; Hanley et al. 2021; Winter et al. 2018; Kunert et al. 2026a, b) or whole leaves (Kunert & Hajek 2022; Visakorpi et al. 2024; Manzi et al. 2025) were put on moist tissues or thin water films in Petri dishes. In some cases, this was done not only for 1 day, but for extended periods of up to 14 days (Winter et al. 2025). Experimental work with detached leaves or leaf disks might already be questioned in the usual procedures of heat exposure experiments when the samples are prepared immediately before the heat treatment and measured right afterwards following dark adaptation (Didion-Gency et al. 2025). Yet keeping detached leaves or leaf disks for days or even weeks after the treatment in the laboratory might be too artificial, leaving it open whether non-recovery or intensified damage would have happened if the leaves were connected to the plant, especially if the plant was well water-supplied. These uncertainties suggest that there is an urgent need for an increased number of thoroughly conducted experiments of heat recovery in trees with more sophisticated methods. It is, so far, unproven that storing leaves or leaf disks in plastic bags or under not specified conditions at dark in the laboratory (Drake et al. 2018; Hanley et al. 2021; Manzi et al. 2025) or in Petri dishes at low light levels (e.g. Perez et al. 2021; Slot et al. 2021; Tiwari et al. 2021; Kunert et al. 2022) represent valid methods to test the capacity for heat recovery in trees under ambient conditions. Therefore, most specifications on reversible versus irreversible PS II heat damage should be regarded with care at the present state of knowledge.

The most comprehensive study of PS II heat stress recovery, we are aware of, has so far been published by Didion-Gency et al. (2025). In the Mediterranean conifers *Cupressus sempervirens* and *Pinus halepensis*, they found no recovery within 4 days after exposure to 45–60°C for 30 min in well-watered seedlings. In *Pinus sylvestris*, *F*_v_/*F*_m_ largely recovered within 4 days from exposure to 45°C, but not from 50–60°C. While these recovery studies were done with intact seedlings, where a branch was sealed in a plastic bag that was exposed to heat in a water bath, the effects of the heat treatments immediately after heat exposure were also compared with results from experiments with detached needles (Didion-Gency et al. 2025). In these comparisons, there was no consistent trends for higher *T*_50_” values in intact well-watered seedlings than in detached leaves, which supports the general validity of the results from the numerous studies with detached leaves.

The problems associated to *T*_50_ and *T*_50_^’^, as well as the limited information on the recovery potential of trees in the field following PS II heat damage were tried to address in recent studies by Hauck et al. (2025, 2026) and Dulamsuren et al. (2026) by avoiding these indices and focusing on the textual presentation of regression models analyzing the effect of temperature, incubation time, and tree species on *F*_v_/*F*_m_. This practice allows an accentuated discussion of relative differences between species within a study, but is hardly suitable for quick comparisons across different studies. Aitken et al. (2026) relied on *F*_v_/*F*_m_ values ≤0.3 as a threshold that was assume to indicative of irreversible PS II damage. This threshold is certainly more likely to be true than the threshold of ≤0.5 that underlies *T*_50_, but nonetheless remains (like *T*_50_) arbitrary, because both thresholds are not supported by any mechanistic explanation that proves that exactly 0.3 or 0.5 are critical *F*_v_/*F*_m_ values.

As researchers obviously see a demand for an index that ranks the heat tolerance of a species in a comparable way to *P*_50_ and *P*_88_ as indicators of drought tolerance (i.e. the water potential at 50% or 88% of embolism in the sapwood cross-section; Delzon & Cochard 2014), we suggest the use of the temperature at the inflection point (here termed *T*_IP_) of the sigmoid regression curve describing the decrease of absolute *F*_v_/*F*_m_ with increasing temperature as a new metric that overcomes most of the expounded problems with *T*_50_ and *T*_50_^’^, except the need to take into account the significance of incubation time. Unlike *T*_50_, *T*_IP_ is not arbitrary, as it represents the temperature of the most rapid decline of *F*_v_/*F*_m_ due to heat exposure. In contrast to *T*_50_^’^, which is based on relative values of *F*_v_/*F*_m_, *T*_IP_ can be directly linked with the *F*_v_/*F*_m_ level, at which the most rapid decline of *F*_v_/*F*_m_ occurs (here termed *F*_IP_), that we consider as an important indicator of species-specific PS II heat tolerance. Existing datasets that were previously analyzed by calculating *T*_50_ or *T_50_*^’^ can easily be used to calculate *T*_IP_ and *F*_IP_ if researchers wish to do so, which is a practical advantage of our new metrics.

We applied *T*_IP_, *T*_50_, and *T*_50_^’^ on data of heat experiments with temperate broadleaved and coniferous tree species and analyzed how these indices responded to incubation time and tree species. We examined whether rankings of tree species according to the PS II heat tolerance differ in dependence of the applied response metric and whether differences in absolute heat tolerance of tree species are suggested depended on the application of *T*_IP_, *T*_50_, or *T*_50_^’^. We assumed that all three parameters are influenced by the duration of heat exposure, but tested the hypothesis that this influence is stronger for the more arbitrary parameter *T*_50_ than for *T*_IP_ and *T*_50_^’^. It is important for us to note that the critical examination of other authors’ methodologies is not meant to lower the high scientific value of the examined papers. All of them contributed to the rapid progress in heat tolerance research and were carried out with valid methods. Rather, with a growing body of literature on this topic, we aim at a critical comparison of methods in order to achieve greater methodological standardization. We focus on heat tolerance research in trees, which is where most recent research has been done due to its significance for climate change ecology, but much of what is said, is also valid for non-woody plants.

## 2. Materials and methods

### 2.1. Examined data sets

We examined existing published (Hauck et al. 2025, 2026) and unpublished datasets of our group, which were all obtained with the same methodology to test the influence of incubation time and to examine differences between *T*_50_, *T*_50_’, and *T*_IP_. All data are from experiments with temperate tree species native or introduced to Central Europe. The published data include data recorded at one point in time from the broadleaved angiosperm trees *Acer campestre* L., *A. platanoides* L., *A. pseudoplatanus* L*., F. sylvatica* L., *Q. petraea* (Matt.) Liebl., *Q. pubescens* Willd., *Q. robur* L., *Q. rubra* L., *Tilia cordata* Mill., and *T. platyphyllos* Scop. and the conifers *Abies alba* Mill., *Picea abies* (L.) H. Karst., *Pinus sylvestris* L., and *Pseudotsuga menziesii* (Mirb.) Franco (Hauck et al. 2025). These were summarized to the comparison of broadleaves versus conifers in Fig. 1. The data in the other figures and tables refer to seasonal species were the same species were exposed to and chlorophyll fluorescence was measured repeatedly from fresh sample collected throughout the growing season of 2025. Species investigated this way included *Fagus sylvatica*, *F. orientalis* Lipsky, *Pseudotsuga menziesii* (data from Hauck et al. 2026) and *Abies alba*, *Larix decidua* Mill., *Picea abies*, and *Pinus sylvestris* (unpublished data). The methodologies of sampling and sample storage are described in Hauck et al. (2025, 2026) and are the same for the so far unpublished data.

**Fig. 1.**
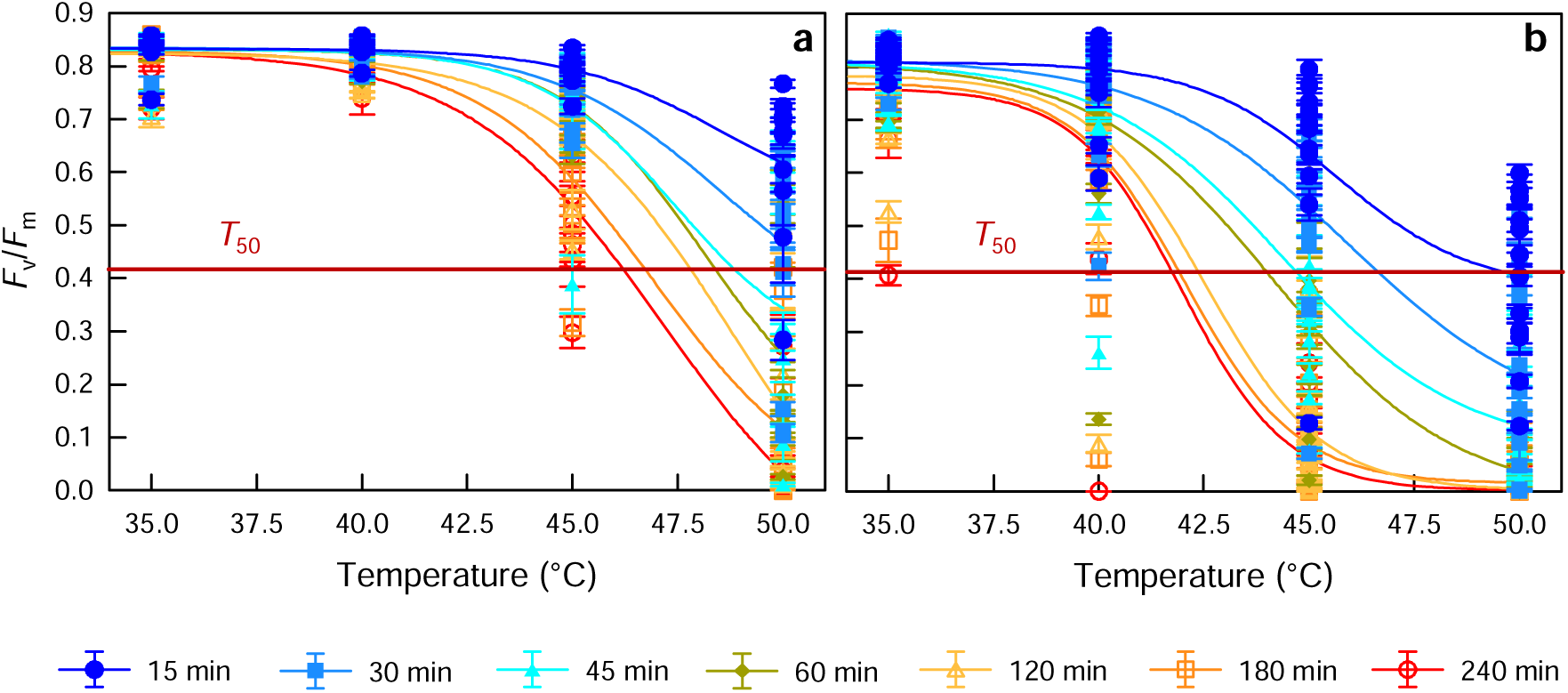
Maximum efficiency at PS II (*F*_v_/*F*_m_) after heat treatments at 35, 40, 45, and 50°C for 15 to 240 min in (a) temperate broadleaved (*N*= 10 species) and (b) coniferous (*N*= 5) tree species from Central Europe. The red horizontal lines mark the level of 50% of maximum *F*_v_/*F*_m_ (*T*_50_).

### 2.2. Experimental setup for heat treatments

The experimental setup in the laboratory is described in detail in Hauck et al. (2025, 2026) and Dulamsuren et al. (2026). We worked with complete detached tree leaves, which were put in paper bags to include some air and then sealed in heat-resistant plastic bags to keep the sample dry. The plastic bags were then incubated in a preheated water bath, after the target temperature had been reached. The interior of the sample bags reached the same temperature as in the water bath, as was controlled with temperature sensors (TC 319, Dostmann electronic, Wertheim, Germany) in control measurements published in Hauck et al. (2025). The temperature was set to either 35, 40, 45, or 50 °C. Incubation time was set to either 15, 30, 45, 60, 120, 180, or 240 min. We used fresh leaves for every combination of temperature and incubation time.

### 2.3. Chlorophyll fluorescence analysis

The paper bags were kept at dark for a minimum of 20 min to allow for dark adaptation. The Mini-PAM-II Photosynthesis Yield Analyzer (Walz, Effeltrich, Germany) was used to measure chlorophyll fluorescence, firstly, with a weak measuring beam to induce electron transport at PS II in order to record minimum fluorescence (*F*_0_). Secondly, we recorded maximum fluorescence (*F*_m_), after the reaction centers of the electron transport chain had been closed by a saturation pulse (5000 µmol m^−2^ s^−1^ for 600 ms). *F*_v_/*F*_m_ was calculated as *F*_v_/*F*_m_ = (*F*_m_ – *F*_0_)*/F*_m_.

### 2.4. Curve fitting and derivation of T_IP_, F_IP_, T_50_, and T_50_’

The relationship *F*_v_/*F*_m_ vs. temperature has a sigmoid shape and can be calculated with several alternative approaches. We calculated *T*_IP_, *F*_IP_, and *T*_50_’ the R package ‘nplr’ 0.1-8 under R 4.4.3. This package applies a 5-parameter model of the generalized logistic function (Richards’ function) that allows for asymmetry of the inflection point and is adjusted to 4 parameters if the curve is symmetric around the inflection point. *T*_IP_ can be extracted from the nplr output as the x-value of the inflection point and *F*_IP_ as the y-value of the inflection point of the regression function. *T*_50_’ can be directly read from the “IC50” value in the nplr output. *R*^2^ values of the models calculated with generalized logistic functions were calculated with nplr to assess goodness of the fits. *T*_50_ was calculated by solving the regression equation to x and setting y to 50% of the maximum *F*_v_/*F*_m_ (i.e. mean *F*_v_/*F*_m_ of the lowest temperature and incubation time). The graphs displayed in Figs. 1 and 2 are based on 4-parameter models with Boltzmann regression, which is a special case of the generalized logistic function and is a good fit for data with a relatively slow transition from the upper to the lower asymptote in comparison with logistic regression. The best fit of the alternative sigmoid functions was tested empirically (Carvalho et al. 2025) and based on this comparison we decided for Boltzmann regression for displaying symmetric curves.

**Fig. 2.**
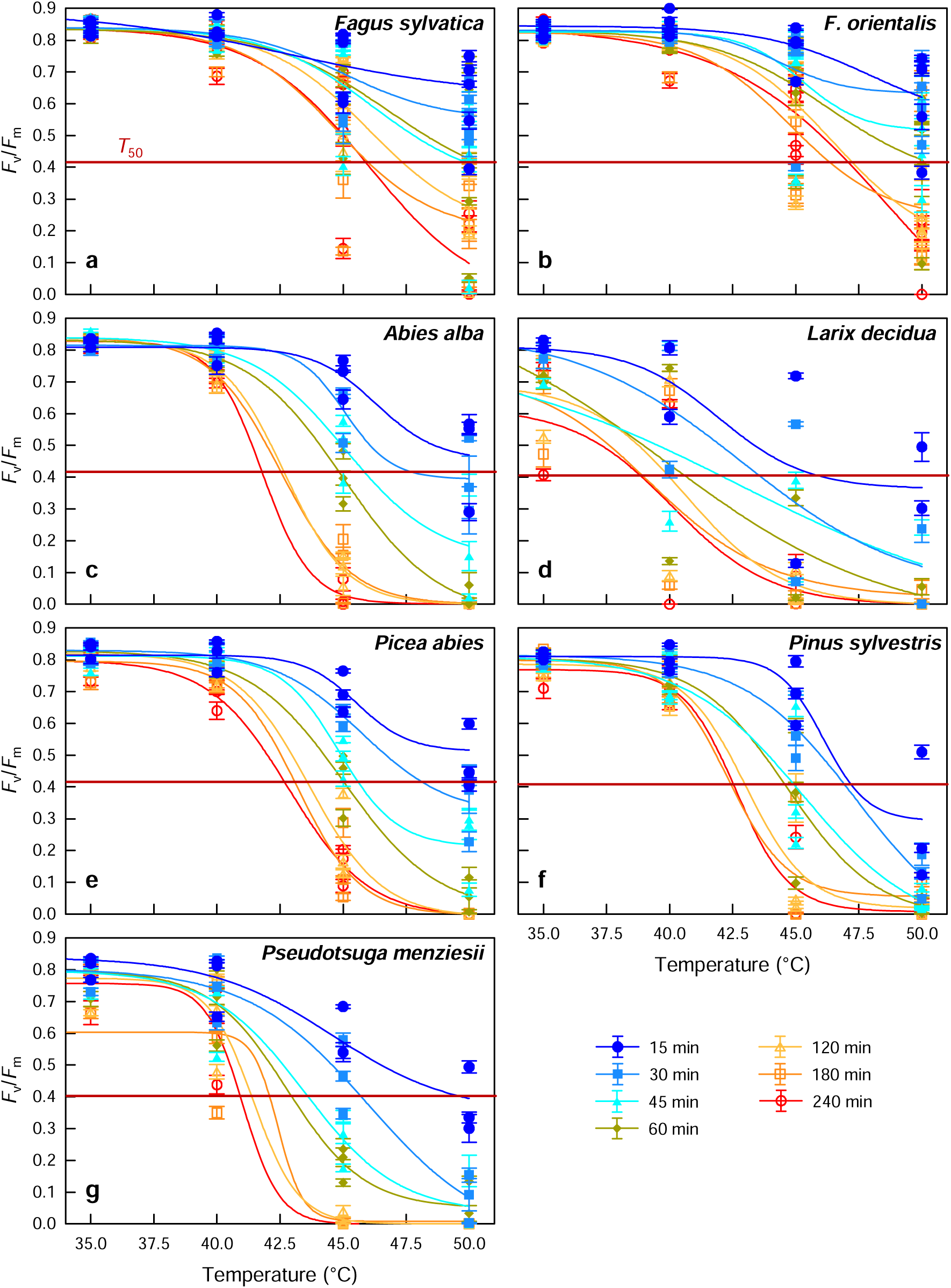
Maximum efficiency at PS II (*F*_v_/*F*_m_) after heat treatments at 35, 40, 45, and 50°C for 15 to 240 min in different tree species. The red horizontal lines mark the level of 50% of maximum *F*_v_/*F*_m_ (*T*_50_): (a) *Fagus sylvatica*, (b) *Fagus orientalis*, (c) *Abies alba*, (d) *Larix decidua*, (e) *Picea abies*, (f) *Pinus sylvestris*, (g) *Pseudotsuga menziesii*.

### 2.5. Statistical analysis

Arithmetic means ± standard errors (SE) are presented throughout the paper. All statistical calculations were computed in R 4.4.3. Ordinary least square linear regression models were used to analyze the influence of temperature, incubation time, and tree species on chlorophyll fluorescence parameters. Interactions could not be analyzed due to multicollinearity, as tested with the R package ‘performance’ 0.17.0. In the analysis of effects of incubation time, *T*_IP_, and *T*_50_, the thermotolerance parameter *T*_50_’ had to be removed from the model due to multicollinearity.

## 3. Results

### 3.1. Time-dependence of F_v_/F_m_ vs. temperature

Incubation time had a strong influence on the response of *F_v_/F_m_* to temperature. Especially short heat treatments of 15 or 30 min had much weaker effects than long treatments of 1–4 h (Fig. 1). Broadleaved angiosperm trees (Fig. 1a) were generally less sensitive to heat than conifers (Fig. 1b). For the mean of broadleaved trees, the regression lines for *F_v_/F_m_* vs. temperature did not reach *T*_50_ (i.e. the line of 50% reduction of *F_v_/F_m_*) at all for samples exposed to temperatures from 35– 50°C for 15 and 30 min (Fig. 1a). For conifers, *T*_50_ was reached at 50.0°C after 15 min and at 46.7°C after 30 min of heat exposure (Fig. 1b). Samples exposed to heat for 1–2 h reached *T*_50_ at a temperature range of 41.8–42.4°C in conifers, but at 46.2–47.9°C in broadleaved trees (Fig. 1).

This general difference between broadleaves and conifers became also evident from the analysis of individual species (Fig. 2; Table 2). Incubation time had a significant negative effect on *T*_50_ in the regression model (Table 2). The broadleaves *Fagus orientalis* and *F. sylvatica* had significantly higher *T*_50_ values than the conifers (Table 2). *T*_50_ was lower in the deciduous conifer *Larix sibirica* than in the evergreen species *Abies alba*, *Picea abies*, *Pinus sylvestris*, and *Pseudotsuga menziesii* (Table 2; Fig. 2). Across the tree species, the goodness of the sigmoid fits increased non-linearly with increasing incubation time and reached a plateau of *R*^2^ = 0.86 to 0.90 at 2–4 h of heat exposure, indicating less variance between the individual curves with at long incubation times (Fig. 3).

**Fig. 3.**
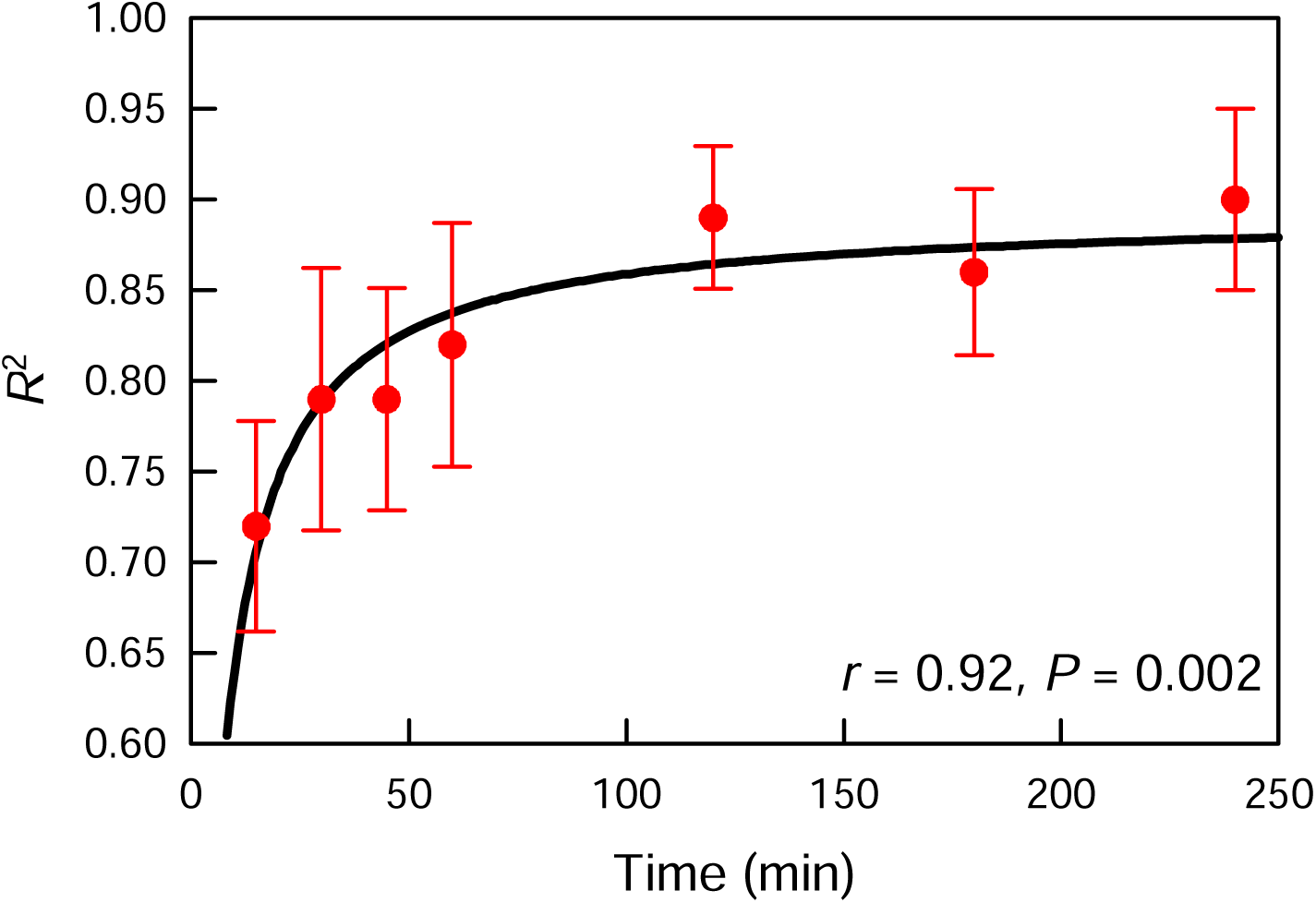
Goodness of the fits measured as *R*^2^ of generalized logistic function calculated after 15 min – 4 h of heat exposure. Means (±SE) of data from *Fagus sylvatica*, *F. orientalis*, *Abies alba*, *Larix decidua*, *Picea abies*, *Pinus sylvestris*, *Pseudotsuga menziesii*.

**Table 2.** Linear regression models analyzing effects of tree species and incubation time on *T*_IP_, *T*_50_, *T*_50_’, and *F*_IP_.

| Parameter | Estimate | SE | P | CI |  |
| --- | --- | --- | --- | --- | --- |
|  |  |  |  | 2.5% | 97.5% |
| $T_{IP}$: | | | | | |
| (Intercept)<sup>a</sup> | 44.30 | 0.46 | <**0.001** | 43.40 | 45.21 |
| Incubation time | −1.18 | 0.19 | <**0.001** | −1.56 | −0.79 |
| *Fagus orientalis* | 2.76 | 0.75 | <**0.001** | 1.24 | 4.28 |
| *Fagus sylvatica* | 1.77 | 0.69 | **0.02** | 0.37 | 3.18 |
| *Picea abies* | −0.12 | 0.66 | 0.86 | −1.45 | 1.21 |
| *Abies alba* | −1.07 | 0.66 | 0.11 | −2.41 | 0.26 |
| *Pseudotsuga menziesii* | −1.38 | 0.63 | **0.04** | −2.66 | −0.10 |
| *Larix decidua* | −3.77 | 0.63 | <**0.001** | −5.04 | −2.49 |
| $R^2$ | 0.79 | | | | |
| $T_{50}$: | | | | | |
| (Intercept)<sup>a</sup> | 44.23 | 0.51 | <**0.001** | 43.19 | 45.28 |
| Incubation time | −1.79 | 0.22 | <**0.001** | −2.23 | −1.35 |
| *Fagus orientalis* | 4.39 | 0.86 | <**0.001** | 2.64 | 6.14 |
| *Fagus sylvatica* | 4.05 | 0.80 | <**0.001** | 2.44 | 5.68 |
| *Picea abies* | 0.49 | 0.76 | 0.52 | −1.05 | 2.02 |
| *Abies alba* | 0.08 | 0.76 | 0.91 | −1.45 | 1.62 |
| *Pseudotsuga menziesii* | −0.75 | 0.72 | 0.31 | −2.22 | 0.73 |
| *Larix decidua* | −2.74 | 0.72 | <**0.001** | −4.21 | −1.27 |
| $R^2$ | 0.82 | | | | |
| $T_{50}'$: | | | | | |
| (Intercept)<sup>a</sup> | 43.77 | 0.70 | <**0.001** | 42.36 | 45.19 |
| Incubation time | −2.02 | 0.27 | <**0.001** | −2.56 | −1.48 |
| *Fagus orientalis* | 4.41 | 0.99 | <**0.001** | 2.41 | 6.40 |
| *Fagus sylvatica* | 4.83 | 0.99 | <**0.001** | 2.83 | 6.82 |
| *Picea abies* | 0.85 | 0.99 | 0.39 | −1.14 | 2.85 |
| *Abies alba* | 0.43 | 0.99 | 0.67 | −1.57 | 2.42 |
| *Pseudotsuga menziesii* | −1.25 | 0.99 | 0.21 | −3.25 | 0.75 |
| *Larix decidua* | −2.02 | 0.27 | <**0.001** | −5.66 | −1.67 |
| $R^2$ | 0.80 | | | | |
| $F_{IP}$: | | | | | |
| (Intercept)<sup>a</sup> | 0.416 | 0.025 | <**0.001** | 0.365 | 0.467 |
| Incubation time | −0.061 | 0.010 | <**0.001** | −0.081 | −0.042 |
| *Fagus orientalis* | 0.158 | 0.036 | <**0.001** | 0.086 | 0.230 |
| *Fagus sylvatica* | 0.164 | 0.036 | <**0.001** | 0.092 | 0.236 |
| *Picea abies* | 0.056 | 0.036 | 0.13 | −0.017 | 0.128 |
| *Abies alba* | 0.096 | 0.036 | **0.01** | 0.024 | 0.168 |
| *Pseudotsuga menziesii* | −0.012 | 0.036 | 0.74 | −0.085 | 0.060 |
| *Larix decidua* | 0.012 | 0.036 | 0.75 | −0.061 | 0.084 |
| $R^2$ | 0.69 | | | | |

<sup>a</sup> *Pinus sylvestris* was selected as baseline species, as $T_{50}$ vs. incubation time diverged least from $T_{IP}$ vs. incubation time in this species (see Fig. 4f).

### 3.2. Differences between T_IP_, T_50_ and T_50_’

*T*_50_ and *T*_50_’ were more strongly influenced by incubation time than *T*_IP_, but for all three parameters this influence was highly significant (Table 2). In several cases, *T*_50_ could not be determined after short heat treatments of less than 1 h, because *F_v_/F_m_* was not reduced to 50% of the start values, which means that *F_v_/F_m_* was above c. 0.4 (Table 3; Fig. 2). This applied to *Abies alba* and *Picea abies* at 15 min, *Fagus sylvatica* at 15–30 min, and *Fagus orientalis* at 15–45 min of heat exposure.

**Table 3.** *T*_IP_, *T*_50_, *T*_50_’, and *F*_IP_ a for selected temperate tree species determined after 15 min and 4 h of heat exposure to 35–50°C.

| Species | $T_{IP}$ | | $T_{50}$ | | $T_{50}'$ | | $F_{IP}$ | |
| --- | --- | --- | --- | --- | --- | --- | --- | --- |
|  | 15 min | 4 h | 15 min | 4 h | 15 min | 4 h | 15 min | 4 h |
| *Fagus orientalis* | 45.4 | 47.0 | n.d.<sup>a</sup> | 47.1 | 50.0 | 45.9 | 0.73 | 0.41 |
| *Fagus sylvatica* | 45.2 | 46.3 | n.d. | 45.9 | 50.0 | 45.0 | 0.72 | 0.38 |
| *Pinus sylvestris* | 47.6 | 42.4 | 47.2 | 42.5 | 47.6 | 42.4 | 0.48 | 0.41 |
| *Abies alba* | 47.0 | 41.6 | n.d. | 41.8 | 49.2 | 41.3 | 0.61 | 0.43 |
| *Picea abies* | 46.6 | 42.8 | n.d. | 42.7 | 49.5 | 41.5 | 0.62 | 0.39 |
| *Pseudotsuga menziesii* | 45.9 | 40.6 | 49.7 | 40.9 | 46.8 | 40.3 | 0.54 | 0.38 |
| *Larix decidua* | 41.2 | 39.8 | 45.8 | 38.8 | 45.8 | 38.8 | 0.61 | 0.33 |

<sup>a</sup> n.d., not determinable for the studied temperature range

**Table 4.** Linear regression model analyzing effects of *T*_IP_, *T*_50_ and incubation time on *F*_IP_.

| Parameter | Estimate | SE | *P* | CI |  |
| --- | --- | --- | --- | --- | --- |
|  |  |  |  | 2.5% | 97.5% |
| (Intercept)<sup>a</sup> | 0.451 | 0.007 | <0.001 | 0.438 | 0.465 |
| $T_{IP}$ | −0.092 | 0.014 | <0.001 | −0.120 | −0.063 |
| $T_{50}$ | 0.131 | 0.015 | <0.001 | 0.101 | 0.161 |
| Incubation time | −0.007 | 0.008 | 0.39 | −0.022 | 0.009 |
| $R^2$ | 0.74 | | | | |

The order of species differed between *T*_IP_, *T*_50_, and *T*_50_’, when they were sorted by heat tolerance. According to linear regression modeling, considering all variants of temperature and incubation time in the same model (Table 2), heat tolerance declined in the order *Fagus orientalis* > *Fagus sylvatica* > *Pinus sylvestris*, *Picea abies*, *Abies alba* > *Pseudotsuga menziesii* > *Larix decidua*, if the assessment was based on *T*_IP_. This ranking was similar for *T*_50_, but heat tolerance of *Pseudotsuga menziesii* did not differ from that of *Pinus sylvestris*, *Picea abies*, and *Abies alba*. Using *T*_50_^’^ instead of *T*_50_ did not result in the detection of a difference in heat tolerance between *Pseudotsuga menziesii*, *Pinus sylvestris*, *Picea abies*, and *Abies alba* either. Moreover, applying *T* ^’^ suggested that *Fagus sylvatica* was more heat-tolerant than *F. orientalis*, which is counterintuitive given the known distribution ranges of these species.

The dependence of *T*_IP_ and the widely used *T*_50_ on the duration of heat exposure differed between tree species and between the two parameters (Fig. 4). In *Pinus sylvestris*, *T*_IP_ and *T*_50_ showed no difference in the response to incubation time (Fig. 4f). In all other species, *T*_IP_ and *T*_50_ attained similar values at long incubation times of 2–4 h, but diverged to varying degrees for short incubation periods (Fig. 4). In general, *T*_50_ tended to be higher than *T*_IP_ at short incubation times, which was most extreme in *Fagus* (Fig. 4a, b) and *Larix* (Fig. 4d), but also evident in the other conifers except *Pinus sylvestris*. *T*_50_ decreased non-linearly with increasing incubation time in all species (Fig. 4). By contrast, there were species-specific differences in the response of *T*_IP_ to incubation time. In *Fagus orientalis* and *F. sylvatica*, there was no significant response of *T*_IP_ to incubation time (Fig. 4a, b). In most other species, *T*_IP_ increased less strongly with decreasing incubation time than *T*_50_.

**Fig. 4.**
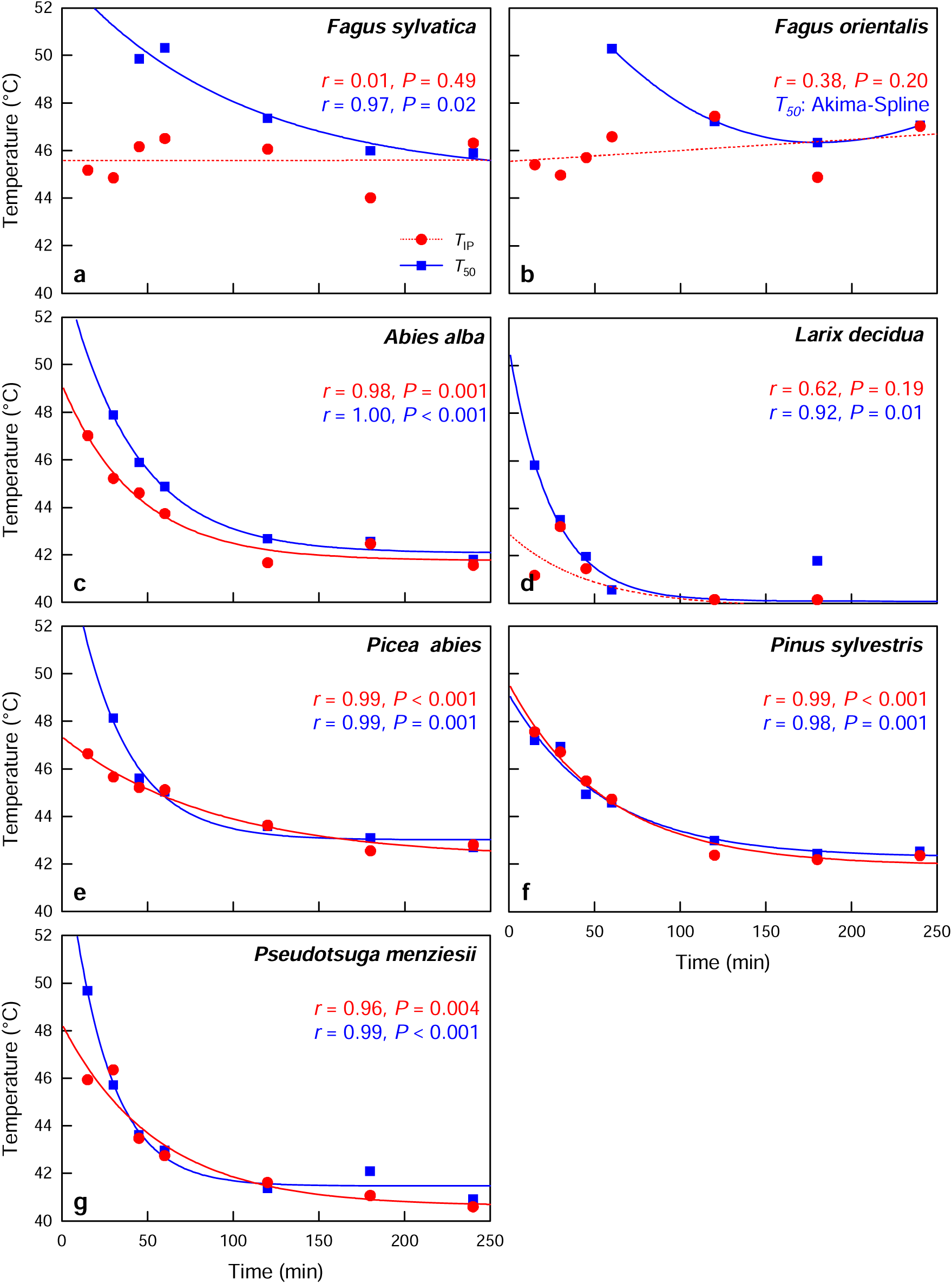
Temperature at inflection point (*T*_IP_) of the regression line for temperature vs. incubation time and temperature of 50% *F*_v_/*F*_m_ compared to maximum *F*_v_/*F*_m_ (*T*_50_) in different tree species: (a) *Fagus sylvatica*, (b) *Fagus orientalis*, (c) *Abies alba*, (d) *Larix decidua*, (e) *Picea abies*, (f) *Pinus sylvestris*, (g) *Pseudotsuga menziesii*.

*T*_IP_ increased linearly with increasing *T*_50_ with a regression slope of 0.71 and intercept of 11.97 (Fig. 5a), which means that at low temperatures *T*_IP_ exceeded *T*_50_, but at high temperatures *T*_IP_ was lower than *T*_50_. With increasing *T*_50_^’^ values, *T*_IP_ increased following a logistic relationship (Fig. 5b). In Fig. 6, we applied the regression functions from Fig. 5 to explore the relationship between *T*_IP_, *T*_50_, and *T*_50_’ based on simulated data. If *T*_50_ was examined relative to *T*_IP_ (Fig. 6a), we found similar results in the narrow temperature range of 41–42°C. Below 41°C, *T*_50_ attained lower values than *T*_IP_, which means that *T*_50_ underestimated heat tolerance relative to *T*_IP_. Above 43°C, *T*_50_ increasingly overestimated heat tolerance relative to *T*_IP_ (Fig. 6a). A similar comparison of *T*_50_’ with *T*_IP_ yielded contrasting results (Fig. 6b): *T*_50_’ overestimated heat tolerance relative to *T*_IP_ at temperatures <46°C, but underestimated heat tolerance at temperatures >46°C. Only at 46°C, results derived from *T*_IP_ and *T*_50_’ were the same (Fig. 6b). A comparison of *T*_50_ and *T*_50_’ showed similar results on in the narrow temperature range of 45–46°C (Fig. 6c). At <45°C*, T*_50_’ overestimated heat tolerance relative to *T*_50_, and at >46°C heat tolerance was underestimated by *T*_50_’ (Fig. 6c). While the difference between *T*_IP_ and *T*_50_ decreased with incubation time in a linear regression model with real data in the studied temperature range of 35–50°C (Table S1), which is consistent with more readings above than below the 100% mark in Fig. 6a with simulated data, there was no such influence of time in a regression model for the difference *T*_IP_ and *T*_50_’ (Table S1).

**Fig. 5.**
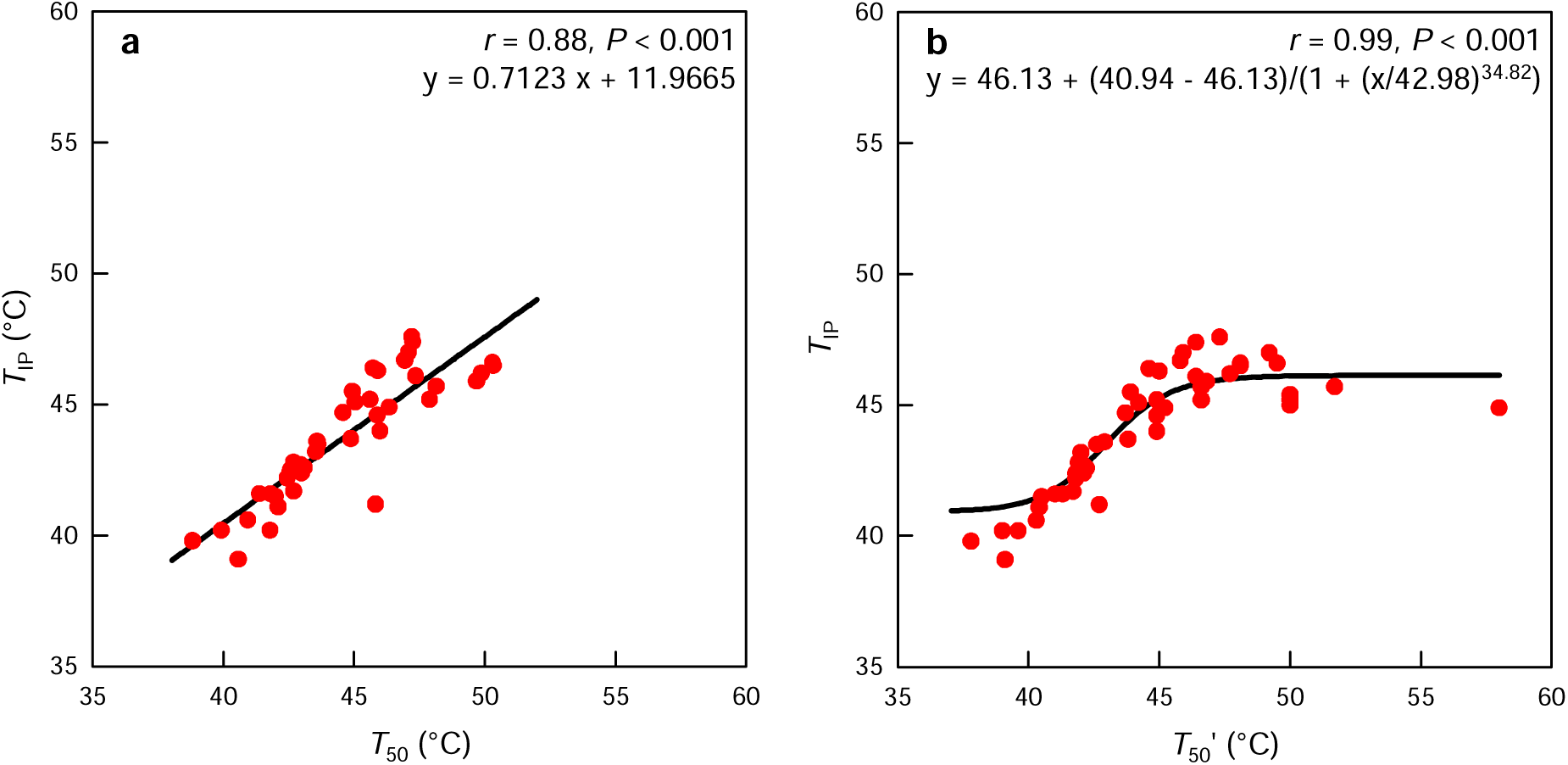
*T*_IP_ versus (a) *T*_50_ and (b) *T*_50_’. Each dot represents a combination of tree species (*Fagus sylvatica*, *F. orientalis*, *Abies alba*, *Larix decidua*, *Picea abies*, *Pinus sylvestris*, *Pseudotsuga menziesii*) with a certain incubation time (15 min – 4 h).

**Fig. 6.**
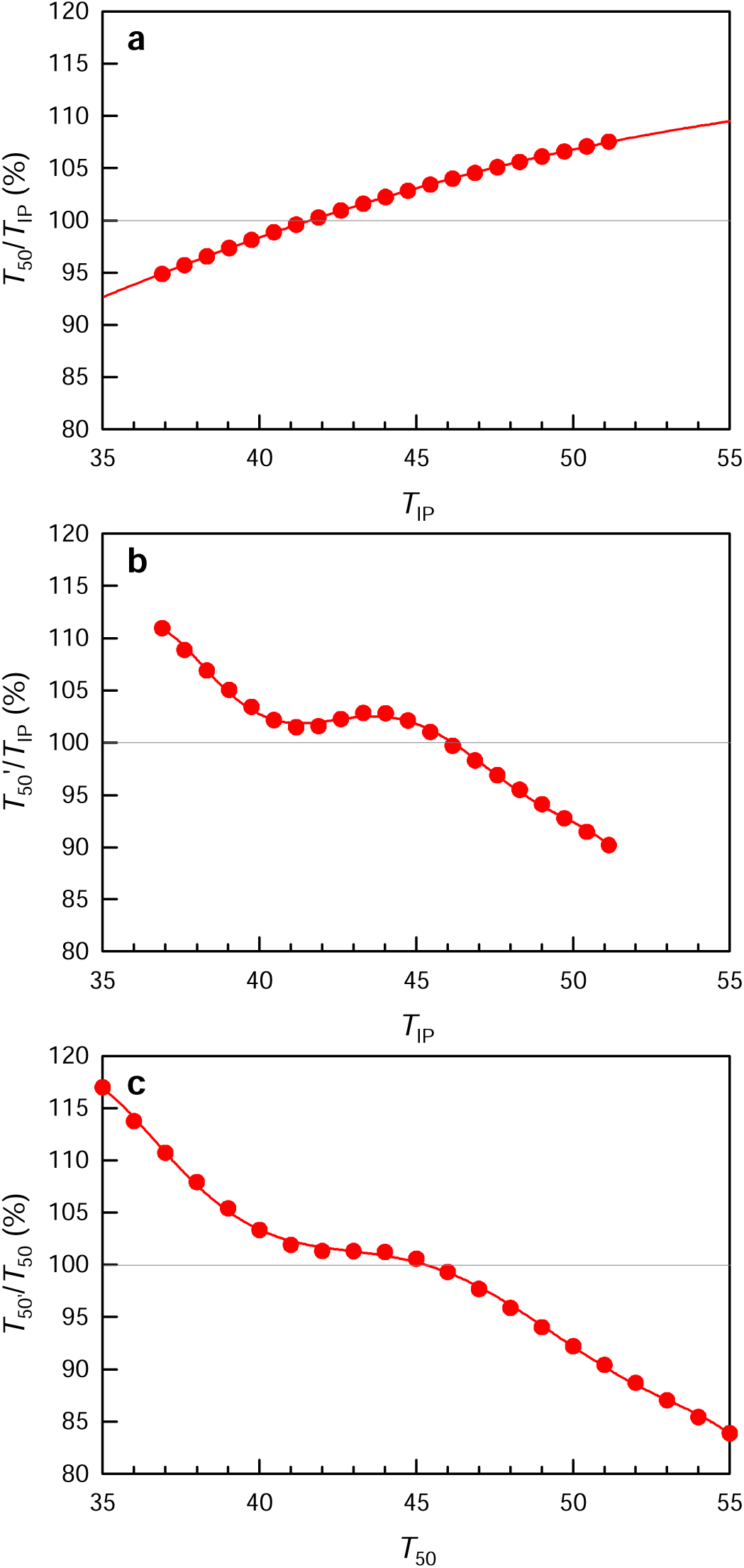
Simulated *T*_IP_, *T*_50_ and *T*_50_’ values based on the regression functions determined in Fig. 5. The vertical axis represents the deviation from the parameter plotted on horizontal axis in percent: (a) *T*_50_ relative to *T*_IP_, (b) *T*_50_’ relative to *T*_IP_, (c) *T*_50_’ relative to *T*_50_. Polynomial regressions of (a) 2^nd^ and (b, c) 7^th^ order. The grey horizontal line represents the curve section where both parameters deliver similar results.

### 3.3. F_v_/F_m_ at T_IP_

*F_IP_* (i.e. *F_v_/F_m_* at *T*_IP_) decreased with increasing incubation time and varied across tree species (Table 2; Fig. 7). In *Pinus sylvestris*, *F*_IP_ was not influenced by incubation time and was close to 0.4 (Fig. 7c), which is consistent with the very similar courses of *T*_IP_ and *T*_50_ versus incubation time in Fig. 4f. In all other tree species, *F*_IP_ decreased with increasing incubation time (Fig. 7). After 4 h of heat exposure, *F*_IP_ came close to 0.4 in most species (Fig. 7), which is consistent with the approximation of *T*_IP_ and *T*_50_ at long incubation times in Fig. 4. In *Larix decidua*, *F*_IP_ was reduced to 0.33 after 4 h of heat exposure (Table 3). Short heat exposure of 15 min reduced *F*_IP_ to 0.54–0.73 in all species except for *Pinus sylvestris* (Table 3). The reduction of *F*_IP_ by 15 min heat exposure was small in *Fagus orientalis* (*F*_IP_ = 0.73) and *F. sylvatica* (*F*_IP_ = 0.72). Reduction was moderate in *Abies alba*, *Larix decidua*, and *Picea abies* (*F*_IP_: 0.61–0.62), but more severe in *Pseudotsuga menziesii* (*F*_IP_ = 0.54) and *Pinus sylvestris* (*F*_IP_ = 0.48).

**Fig. 7.**
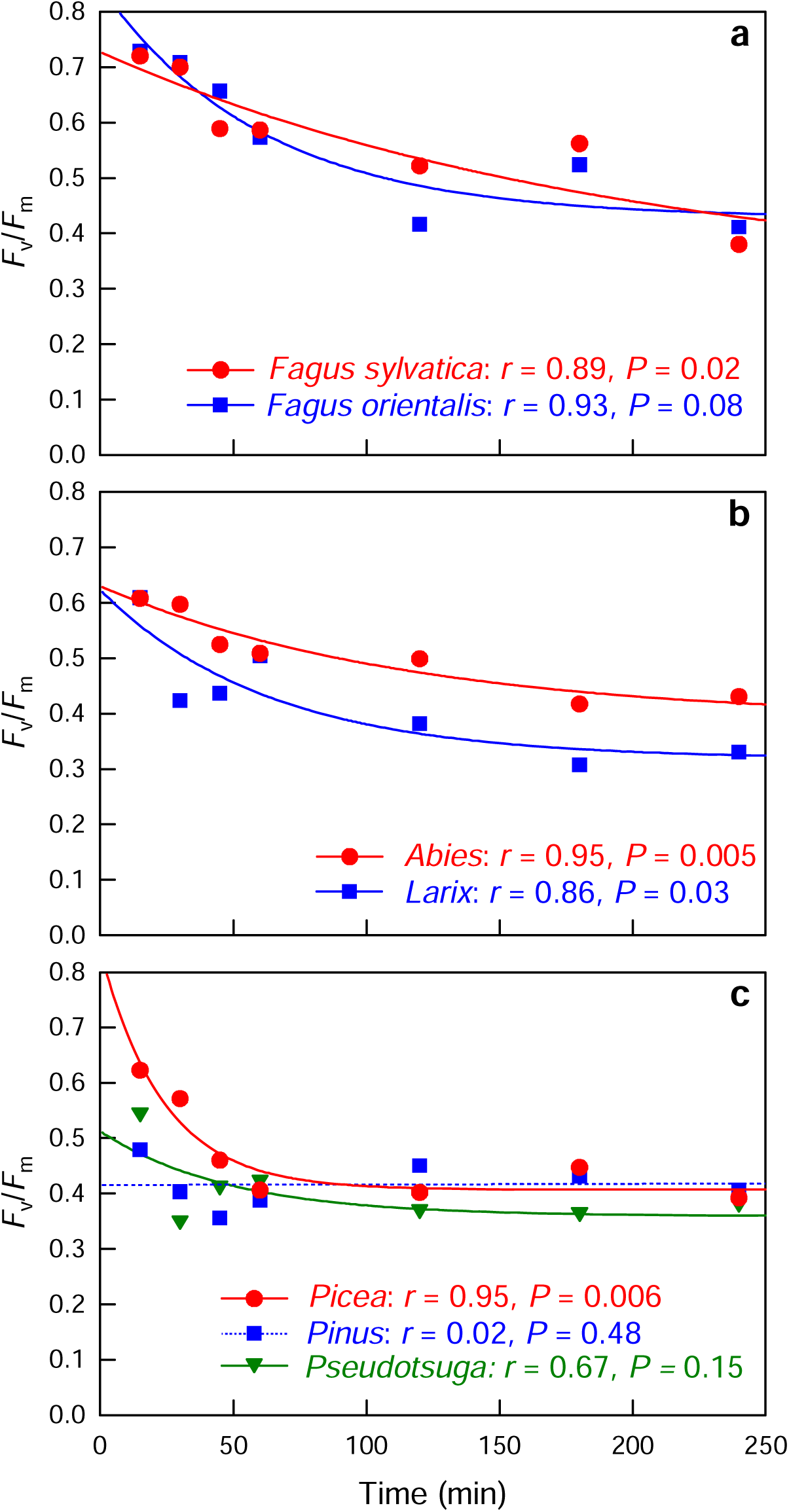
F_IP_ (i.e. maximum quantum yield [*F*_v_/*F*_m_] at *T*_IP_) in different tree species: (a) *Fagus sylvatica*, *Fagus orientalis*; (b) *Abies alba*, *Larix decidua*; (c) *Picea abies*, *Pinus sylvestris*, *Pseudotsuga menziesii*.

## 4. Discussion

### 4.1. Time-dependence of PS II heat stress

Our analysis showed a significant negative effect of the duration of the heat treatment in the experiment on the different parameters (*T*_IP_, *T*_50_, *T*_50_’) that are supposed to characterize the temperature at which PS II is assumed to suffer from critical damage. The same was true for *F*_IP_, representing *F_v_/F_m_* at *T*_IP_. These results confirm the high significance of incubation time for the results of heat tolerance experiments in trees (and other plants) that was already pointed out by Neuner & Buchner (2023) and Hauck et al. (2025) and leads to the conclusion that specifications of temperatures, which are thought to be critical for tree survival, are only comparable across different studies if incubation times is kept constant. Our data (e.g. Figs. 2, 4) suggest that even varying the incubation time between 15 and 30 min leads to very different results in terms of *T*_IP_, *T*_50_, and *T*_50_’ values. For *T*_50_” the difference between 15 and 30 min was already substantiated by Didion-Gency et al. (2025), but most studies using incubation times of either 15 or 30 min (Table 1) treated results obtained with 15 or 30 min as comparable. Our data show that the results stabilize at long incubation times of 2–4 h, which is suggestive of more strongly focusing on such timespans in future experiments. Such incubation times seem also to be more realistic with respect to ambient conditions, as heat waves intensify over time with progressive drying of soils and vegetation on the landscape scale, when the mitigating effect of latent heat on atmospheric warming is increasingly diminished (Miralles et al. 2014). The recent heat wave of 2026 in Western and Central Europe, when maximum air temperatures of 41.8°C in Germany and 43.8°C in France were reached (WMO 2026), is an outstanding example of this. Canopy temperatures usually even exceed air temperatures (Still et al. 2022; Endris & Rehm 2025; Dulamsuren et al. 2026). Regression modeling suggested a greater negative effect of incubation time on *T*_50_ and *T*_50_’ than on *T*_IP_, which partly confirms our corresponding hypothesis and suggests that *T*_IP_ is more stable to variations in the duration of heat exposure than the two other parameters.

### 4.2. Explanatory power of T_50_, T_50_’, and T_IP_ to characterize heat tolerance in tree species

An important message of this paper is that *T*_50_, *T*_50_’, and *T*_IP_ cannot be used interchangeably, because they can provide very different results for the same tree species under the same heat treatment. While *T*_IP_ is novel and for the first time defined in this paper, *T*_50_ and *T*_50_’ have been used repeatedly (studies cited in Table 1). These parameters were defined properly in all individual studies, but have never been discussed as to differ fundamentally from each other before. Because we could show that there is this difference between *T*_50_ and *T*_50_’, it does not make much sense to continue using both parameters without specifically interpreting them in future studies. Rather, we suggest to unify parametrization and to decide for one of these parameters or, as we think, better for the third (novel) parameter *T*_IP_. Since all previous authors have properly defined their parameters, the intention of this paper is not to criticize previous work, but to improve the basis for such work under an increased interest for heat tolerance in trees in the age of climate change. This is because our analyses substantiate that the definition of heat tolerance parameters matters a lot for the assessment of the heat tolerance of individual species and as therefore the differences, as well as advantages and disadvantages of the individual parameters should be discussed.

Different absolute temperatures that can be assessed as critical for tree survival for the same tree species dependent on the use of *T*_50_, *T*_50_’, or *T*_IP_. Furthermore, differences can occur in the order of tree species if sorted by heat tolerance according to *T*_50_, *T*_50_’, and *T*_IP_. These differences can lead to very different assessments as to whether a tree species is considered as heat-tolerant under a given climate or not, influencing silvicultural decision-making. For example, *Pseudotsuga menziesii* has been considered as a very important replacement tree species for the climate change adaptation of Europe’s temperate forests, as it was considered to be more drought-tolerant than widespread native European conifers (Spiecker et al. 2019; Thomas et al. 2022). In terms of heat tolerance, assessments diverge. Based on the discussion of regression models based on experimental heat treatments at 35–50°C for 15 min to 4 h, Hauck et al. (2025, 2026) identified *P. menziesii* as a heat-sensitive temperate tree species, which is not recommendable for climate change adaptation of forestry. In Germany, this recommendation was directly acknowledged on a policy level (Kätzel et al. 2026) compared to recommendations of the Scientific Advisory Boards of Agricultural and Forest Policy (2016), where *P. menziesii* was considered as a suitable candidate to replace *Picea abies* and *Pinus sylvestris* at a large scale and where also the partial replacement of *Fagus sylvatica* by *P. menziesii* has been proposed. In agreement with the detailed discussions of regression models analyzing PS II heat tolerance in response to temperature, incubation time, and tree species by Hauck et al. (2025, 2026), our results based on *T*_IP_ and *F*_IP_ clearly discourage from using *P. menziesii* as a tree species for silvicultural climate change adaptation under a hotter and drier climate. By contrast, results based on *T*_50_ and *T*_50_’ would (though not directly promoting it) not speak against a widespread introduction of *P. menziesii* to adapt Europe’s temperate forests to climate change. The practical significance of these different assessments is obvious and claims for a more detailed discussion of *T*_50_, *T*_50_’, and *T*_IP_. Higher heat tolerance of the more southernly distributed *Fagus orientalis* than of the cool-temperate *Fagus sylvatica* according to *T*_50_ and *T*_IP_, but lower heat tolerance of *F. orientalis* compared to *F. sylvatica* according to *T*_50_’ corroborates the need for a detailed discussion of the explanatory power of these parameters, as also *F. orientalis* is discussed as a replacement tree species for the native *F. sylvatica* in Central Europe under climate change (Budde et al. 2023).

*T*_50_ is most widely used in the literature (Table 1), but has the disadvantage that it represents an arbitrary threshold without mechanistic basis. The same is true for the threshold of *F*_v_/*F*_m_ = 0.3 used by Aitken et al. (2026). It is obvious that the likelihood that heat damage becomes lethal increases, the lower the *F*_v_/*F*_m_ threshold is set. Nevertheless, in practice, defining this threshold meets a lack of sufficient information on critical temperatures for irreversible versus reversible heat tolerance (Didion-Gency et al. 2025). At the current state of knowledge this might be the most critical gap in heat tolerance research.

As already pointed out in the introduction, there is much room for improvement in experiments on recovery from PS II heat damage in trees under realistic conditions. Winter et al. (2025), for instance, pointed out that heat-induced leaf necrosis would be more tightly related to *T*_50_ values recorded 2 weeks after experimental heat exposure than to *T*_50_ determined the day after heat exposure. At first glance, this result points to a lack of recovery and a low explanatory power of *T*_50_ determined shortly after the heat treatment. However, necrosis was not quantified at the time of the first chlorophyll fluorescence analysis the day after the heat treatment (Winter et al. 2025). Therefore, it is not surprising that leaf necrosis (determined 2 weeks later) was more closely related to *T*_50_ at that time than to *T*_50_ the day after the experiment. Since these data are based on leaf disks kept for 2 weeks in the growth chamber, the values after 2 weeks could be challenged, because the conditions in the experiment were obviously very different from heat-exposed tree leaves connected to intact branches under cool conditions and good water supply after the end of a heat wave under ambient conditions. Most other studies of heat stress recovery in trees refer to short recovery during 1 day and were usually also conducted under rather artificial conditions (e.g. Drake et al. 2018; Hanley et al. 2021; Manzi et al. 2025). As long as information about heat recovery of PS II damage remains incomplete, the explanatory power of *T*_50_ is limited. For short incubation times and realistic temperature ranges, *T*_50_ cannot even be determined (Table 3; Fig. 2), which limits its application, as heat treatments to temperatures of even 60 or 70°C are needed to calculate *T*_50_ (Araújo et al. 2021; Perez et al. 2021; Okubo et al. 2023; da Silva & Rossato 2024). These temperature are extremely unrealistic even under the most pessimistic climate change scenarios.

*T*_50_’ avoids this problem by standardization of *F*_v_/*F*_m_ from 0–100% for the lowest and highest observed values, but completely lacks information on the level of *F*_v_/*F*_m_, to which *F*_v_/*F*_m_ is reduced by the heat treatment. In our data, absolute *F*_v_/*F*_m_ values were partly only weakly reduced after heat exposure at short incubation times, like, for example, in *Fagus orientalis*, *F. sylvatica* and also in *Abies alba* (Fig. 2). During standardization of *F*_v_/*F*_m_ while calculating *T*_50_’, the absolute change in *F*_v_/*F*_m_, which is important for the assessment of damage severity and its potential irreversibility is left unattended. For example, *T*_50_’ values of 50.0°C after 15 min heat exposure in both *F. orientalis* and *F. sylvatica* (Table 3) corresponded to *F*_v_/*F*_m_ values of 0.62±0.07 in *F. orientalis* and 0.61±0.06 in *F. sylvatica* after 15 min of 50°C in our data. Even though much more research on reversibility is needed, it seems unlikely that the reduction of *F*_v_/*F*_m_ from ca. 0.8 to 0.6 (i.e. by roughly 25% and thus exceeding the 15%-threshold of *T*_crit_) was irreversible or extremely critical (Kreslavski et al. 2008; Didion-Gency et al. 2025; Hauck et al. 2026). While the specification of *T*_50_’ does not include any information on the severity of *F*_v_/*F*_m_ reduction and thus PS II damage, our novel metric with *T*_IP_ and *F*_IP_ allows to differentiate between different severities of *F*_v_/*F*_m_ declines at the temperature, where the reduction is strongest. For the case example of *F. orientalis* and *F. sylvatica*, *T*_IP_ was determined as, respectively, 45.4 and 45.2°C suggesting critical damage at rather low leaf temperate, whereas the corresponding *F*_IP_ values of 0.73 and 0.72 showed that the degree of *F*_v_/*F*_m_ reduction at this temperature of most rapid reduction in *F*_v_/*F*_m_ was minor, indicating that *F. orientalis* and *F. sylvatica* are both relatively heat-tolerant (Table 3), which is line with results of Hauck et al. (2025, 2026).

Though not explicitly stated in the publications using *T*_50_’ (e.g. Marias et al. 2017; Kunert et al. 2022), this parameter is derived from or has at least parallels to the relative inhibitory concentration (IC_50_) used in pharmacology (Nevozhay 2014). However, it should be considered that pharmacology and PS II tolerance have different backgrounds. In pharmacology, the situation is different, as doses of the medication often have to be limited due to side effects or as the drug under study is simply only partially effective and used in the lack of a more efficient one. In research on the thermal tolerance of plants, it is evident that there is always some treatment that will destroy PS II, which does not seem to make the use of a relative parameter like *T*_50_’ necessary. The only advantage of *T*_50_’ compared to *T*_50_ is that a value can always be calculated, including for experiments using short incubation times in highly heat-tolerant species.

The novel parameters *T*_IP_ and *F*_IP_, which are proposed in this paper have the advantage that their definition is not arbitrary, as in the case of *T*_50_, because the inflection point of a sigmoid relationship marks the curve section, where the strongest changes occur that must be driven by a mechanism, regardless whether the mechanism has been elucidated or not (Goshu & Koya 2013). Calculation the inflection point from absolute values of *F*_v_/*F*_m_ avoids shifts in the critical temperature due to changes in curve shape during standardization. The relevance of this point becomes evident, when *T*_IP_ and *T*_50_’ are compared in Table 3. Moreover, the combination of *T*_IP_ with *F*_IP_, which cannot be determined from standardized curves used for calculating *T*_50_’, allows the specification of the severity of *F*_v_/*F*_m_ decline in the temperature range, where the heat effect on PS II functioning is strongest.

As shown in Fig. 6, *F*_v_/*F*_m_ standardization between 0 and 100% leads to strong differences of *T*_50_’ to *T*_IP_ and *T*_50_ at the lower and the upper ends of the examined temperature range. This limits the comparability of *T*_50_’ to *T*_IP_ and *T*_50_, but nonetheless values of *T*_50_’ and *T*_50_ were, so far, used interchangeably in the literature, which is a practice that should no longer been followed. Using *T*_50_’, Kunert et al. (2026b) concluded in a study of three Mediterranean *Quercus* species that long-term heat exposure >1 h would blur existing species-specific differences in PS II heat tolerance that were detectable by short-term heat exposure of 15 and 30 min. However, the strong deviation of *T*_50_’ from both *T*_IP_ and *T*_50_ at the lower and the higher ends of the investigated temperature range in Fig. 6b, c and the fact that short-term incubation is likely to produce preferentially higher parameter estimates suggests that some of the estimated variation between species after short incubation might also be the result of an artefact due to using *T*_50_’

## 5. Conclusions

PS II heat tolerance has received increased attention in the recent past and has resulted in a rapidly increasing number of published studies. We could confirm the importance of incubation time in heat exposure experiments for the results and show that *F*_v_/*F*_m_ values stabilize and become increasingly comparable at long incubation time >1 h. We could also show that the multitude of papers has generated deviating parameters of heat tolerance and we argue that the inflection point of the sigmoid *F*_v_/*F*_m_-vs.-temperature relationship (proposed as *T*_IP_) and the corresponding *F*_v_/*F*_m_ value (termed *F*_IP_) provide better substantiated estimates of heat tolerance than *T*_50_ and its derivatives that have been calculated in different ways and are either not linked to a mechanism that would suggest a threshold for critical damage at the specific temperature (*T*_50_) or have less informative value (*T*_50_’, *T*_50_”) than *T*_IP_ in combination with *F*_IP_.

Our paper has been written as a suggestion for standardization and improvement of heat tolerance indices in trees, but does not deny the importance and novelty of the previous studies on heat tolerance, some of which were seminal for the understanding PS II and whole-plant heat tolerance. Our discussion has been exemplified for trees, because the study of the climate change response of forests is particularly important, but we are confident that most of our conclusions are valid for vascular plants in general.

## Acknowledgements

Unpublished data included in this paper were measured in the authors’ work group at the University of Freiburg. Germar Csapek, Klara Krämer, Yves Lucas, Lena Popp, Ole Schmidt, and Linda Szafranek is thanked for their work in the experiments.

---

**Table S1.**
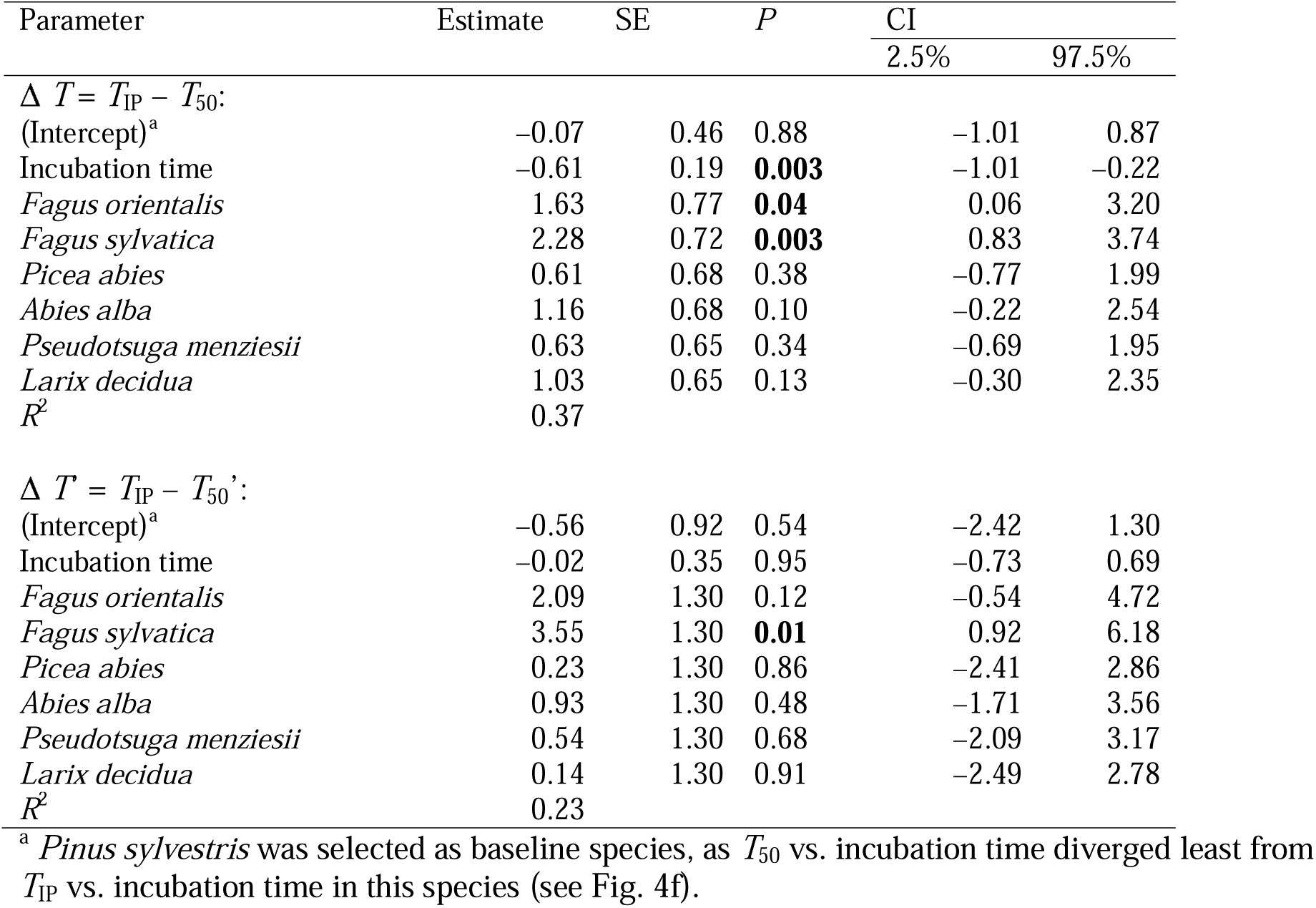
Linear regression models analyzing effects of tree species and incubation time on the difference between both *T*_IP_ and *T*_50_ (Δ *T* = *T*_IP_ – *T*_50_) as well as *T*_IP_ and *T*_50_ (Δ *T*’ = *T*_IP_ – *T*_50_’).

## References

Adhikari Y, Blumröder JS, Meier C, Ibisch PL (2025) Beech buffers: Microclimate regulation in temperate old-growth forests, surroundings and forest edge. Ecol Indic 178:114111

Aitken SM, Arnold PA, Brookhouse MT, Cook AM, Danzey LM, Harris RJ, Leigh A, Nicotra AB (2026) Morphological and heat-tolerance traits are associated with progression and impact of, but not vulnerability to, tree decline. For Ecol Manag 605:123523

Allakhverdiev SI, Kreslavski VD, Klimov VV, Los DA, Carpentier R, Mohanty P (2008) Heat stress: an overview of molecular responses in photosynthesis. Photosynth Res 98:541–550

Araújo I, Marimon BS, Scalon MC, Fauset S, Marimon Junior BH, Tiwari R, Galbraith DR, Gloor MU (2021) Trees at the Amazonia-Cerrado transition are approaching high temperature thresholds. Environ Res Lett 16:034047

Becker FN, Fink AH, Bissolli P, Pinto JG (2022) Towards a more comprehensive assessment of the intensity of historical European heat waves (1979–2019). Atmos Sci Lett 23:e1120

Budde KB, Hötzel S, Müller M, Samsonidze N, Papageorgiou AC, Gailing O (2023) Bidirectional gene flow between *Fagus sylvatica* L. and *F. orientalis* Lipsky despite strong genetic divergence. For Ecol Manag 537:120947

Carvalho FS, Rezende BR, Santos ARPd, Sartori MMP (2025) Modeling seed longevity and percentile prediction: a sigmoidal function approach in soybean, maize, and tomato. AgriEngineering 7:5

Da Silva, B. H. P., & Rossatto, D. R. (2022). Leaves of neotropical savanna tree species are more heat-tolerant than leaves of semi-deciduous forest species. Theoret Exp Plant Physiol 34:227–234

Da Silva BHP, Rossatto DR (2024) Leaf tolerance to heat is independent of leaf phenology in neotropical savanna trees. Trees 38:1343–1350

Dascaliuc A, Ralea T, Cuza P (2007) Influence of heat shock on chlorophyll fluorescence of white oak (*Quercus pubescens* Willd.) leaves. Photosynthetica 45:469–471

Delzon S, Cochard H (2014) Recent advances in tree hydraulics highlight the ecological significance of the hydraulic safety margin. New Phytol 203:355–358

Didion-Gency M, Gauthey A, Johnson KM, Schuler P, Grossiord C (2025) Leaf excision and exposure duration alter the estimates of the irreversible photosynthetic thermal thresholds. Plant Cell Environ 48:5357–5368

Doughty CE, Keany JM, Wiebe BC et al. (2023) Tropical forests are approaching critical temperature thresholds. Nature 621:105–111

Drake JE, Tjoelker MG, Vårhammar A et al. (2018) Trees tolerate an extreme heatwave via sustained transpirational cooling and increased leaf thermal tolerance. Glob Change Biol 24:2390–2402

Dulamsuren C, Byamba-Yondon G, Oyungerel S, Nitschke R, Gebauer T (2023) Non-structural carbohydrate concentrations in contrasting dry and wet years in early- and late-successional boreal forest trees. Trees 37:1315–1332

Dulamsuren C, Abbas JT, Csapek G, Naranbayar E, Uitumen T, Amarjargal D, Byamba-Yondon G, Saindovdon D, Munkhzul T, Batsaikhan G, Hauck M (2026) Heat tolerance and canopy temperatures of Larix sibirica under highly continental climate in Mongolia’s boreal forest. bioRxiv 2026.06.30.735460. 10.64898/2026.06.30.735460

Endris J, Rehm E (2025) Leaf temperatures exceed thermal heat tolerances for a community of eastern North America hardwood trees. AoB Plants 17:plae060

Feeley K, Martinez-Villa J, Perez T, Silva Duque A, Triviño Gonzalez D, Duque A (2020) The thermal tolerances, distributions, and performances of tropical montane tree species. Front For Glob Change 3:25

Goshu AT, Toya PR (2013) Derivation of inflection points of nonlinear regression curves- implications to statistics. Am J Theor Appl Stat 2:268–272

Groover A, Holbrook NM, Polle A et al. (2025) Tree drought physiology: critical research questions and strategies for mitigating climate change effects on forests. New Phytol 245:1817–1832

Guadagno CR, Ewers BE, Speckman HN, Aston TL, Huhn BJ, DeVore SB, Ladwig JT, Strawn RN, Weinig C (2017) Dead or alive? Using membrane failure and chlorophyll *a* fluorescence to predict plant mortality from drought. Plant Physiol 175:223–234

Hanley PA, Arndt SK, Livesley SJ, Szota C (2021) Relating the climate envelopes of urban tree species to their drought and thermal tolerance. Sci Total Enviro 753: 142012

Hauck M, Schneider T, Bahlinger S, Fischbach J, Oswald G, Csapek G, Dulamsuren C (2025) Heat tolerance of temperate tree species from Central Europe. For Ecol Manag 580:122541

Hauck M, Csapek G, Krämer K, Schmidt O, Lucas Y, Popp L, Szafranek L, Dulamsuren C (2026) Heat tolerance and its seasonal acclimation in *Fagus sylvatica* compared to *Fagus orientalis* and *Pseudotsuga menziesii*. bioRxiv 2026.05.17.725742. 10.64898/2026.05.17.725742

Havaux M, Greppin H, Strasser RJ (1991) Functioning of photosystems I and II in pea leaves exposed to heat stress in the presence or absence of light: analysis using in-vivo fluorescence, absorbance, oxygen and photoacoustic measurements. Planta 186:88–98

Húdoková H, Petrik P, Petek-Petrik A, Konôpková A, Leštianka A, Střelcová K, Kmet J, Kurjak D (2022) Heat-stress response of photosystem II in five ecologically important tree species of European temperate forests. Biologia 77:671–680

Hüve K, Bichele I, Rasulov B, Niinemets Ü (2011) When it is too hot for photosynthesis: heat- induced instability of photosynthesis in relation to respiratory burst, cell permeability changes and H_2_O_2_ formation. Plant Cell Environ 34:113–126

Kätzel R, Bauhus J, Endres E et al. (2026) Anpassung von Wäldern durch Unterstützte Migration: Potenziale und Risiken. Scientific Advisory Board of Forest Policy, Federal Ministry of Agriculture, Nutrition, and Homeland. Berlin, Germany (in German)

Konôpková A, Kurjak D, Kmet J, Klumpp R, Longauer R, Ditmarová L, Gömöry D (2018) Differences in photochemistry and response to heat stress between silver fir (*Abies alba* Mill.) provenances. Trees 32:73–86

Krause GH, Winter K, Krause B, Jahns P, García M, Aranda J, Virgo A (2010) High-temperature tolerance of a tropical tree, *Ficus insipida*: methodological reassessment and climate change considerations. Funct Plant Biol 37:890–900

Krause GH, Cheesman AW, Winter K, Krause B, Virgo A (2013) Thermal tolerance, net CO2 exchange and growth of a tropical tree species, *Ficus insipida*, cultivated at elevated daytime and nighttime temperatures. J Plant Physiol 170:822–827

Kunert N, Hajek P (2022) Shade-tolerant temperate broad-leaved trees are more sensitive to thermal stress than light-demanding species during a moderate heatwave. Trees For People 9:100282

Kunert N, Hajek P, Hietz P, Morris H, Rosner S, Tholen D (2022) Summer temperatures reach the thermal tolerance threshold of photosynthetic decline in temperate conifers. Plant Biol 24:1254–1261

Kunert N, Ehrmann J, Gebhard S, Hofmann S, Zimmermann G, Hajek P (2026a) Temperate tree species show cross-tolerance to heat, drought, and late spring-frost stress. New Phytol. 10.1111/nph.71277

Kunert N, Düsterhöft E, Stumpf B (2026b) Heat treatment duration affects in vitro-induced photosynthetic impairment and development of necrotic leaf tissue in three Mediterranean oak species. Plant Biol 28:1256–1267

Kurjak D, Konôpková A, Kmet J, Macková M, Frýdl J, Živčák M, Palmroth S, Ditmarová L, Gömöry D (2019) Variation in the performance and thermostability of photosystem II in European beech (*Fagus sylvatica* L.) provenances is influenced more by acclimation than by adaptation. Eur J Forest Res 138:79–92

Leuzinger S, Körner C (2007) Tree species diversity affects canopy leaf temperatures in a mature temperate forest. Agric For Meteorol 146:29–27

Li X, Wen Y, Chen X, Qie Y, Cao KF, Wee AKS. Correlations between photosynthetic heat tolerance and leaf anatomy and climatic niche in Asian mangrove trees. Plant Biol 24:960–966

Manzi OJL, Mujawamariya M, Tarvainen L, Ziegler C, Andersson MX, Dusenge ME, Fridell A, Reese H, Spetea C, Uwizeye FK, Wittemann M, Nsabimana D, Wallin G, Uddling J (2025) Photosynthetic heat tolerance partially acclimates to growth temperature in tropical montane tree species. Plant Cell Environ 48:7848–7861

Marias DE, Meinzer FC, Woodruff DR, McCulloh KA (2017) Thermotolerance and heat stress responses of Douglas-fir and ponderosa pine seedling populations from contrasting climates. Tree Physiol 37:301–315

Miralles DG, Teuling AJ, van Heerwaarden CC, Vilà-Guerau de Arellano (2014) Mega-heatwave temperatures due to combined soil desiccation and atmospheric heat accumulation. Nature Geosci 7:345–349

Mohanty P, Kreslavski VD, Klimov VV, Los DA, Mimuro M, Carpentier R, Allakhverdiev SI (2012) Heat stress: susceptibility, recovery and regulation. In: Eaton-Rye JJ, Tripathy BC, Sharkey TD (eds) Photosynthesis. Plastid biology, energy conversion and carbon assimilation. Springer, Dordrecht, pp 251–274

Münchinger I, Hajek P, Akdogan B, Caicoya AT, Kunert N (2023) Leaf thermal tolerance and sensitivity of temperate tree species are correlated with leaf physiological and functional drought resistance traits. J For Res 34:63–76

Neuner G, Buchner O (2023) The dose makes the poison: The longer the heat lasts, the lower the temperature for functional impairment and damage. Environ Exp Bot 212:105395

Nevozhay D (2014) Cheburator software for automatically calculating drug inhibitory concentrations from *in vitro* screening assays. PLoS ONE 9:e106186

O’Brien MJ, Leuzinger S, Philipson CD, Tay J, Hector A (2014) Drought survival of tropical tree seedlings enhanced by non-structural carbohydrate levels. Nat Clim Change 4:710–714

Okubo N, Inoue S, Ishii HR (2023) Tolerance and acclimation of the leaves of nine urban tree species to high temperatures. Forests 14:1639

Perez TM, Feeley KJ (2020) Photosynthetic heat tolerances and extreme leaf temperatures. Funct Ecol 34:2236–2245

Perez TM, Socha A, Tserej O, Feeley KJ (2021). Photosystem II heat tolerances characterize thermal generalists and the upper limit of carbon assimilation. Plant Cell Environ 44:2321–2330

Petrik P, Petek-Petrik A, Konôpková A, Fleischer P, Stojnic S, Zavadilova I, Kurjak D (2023) Seasonality of PSII thermostability and water use efficiency of in situ mountainous Norway spruce (*Picea abies*). J For Res 34:197–208

Scientific Advisory Boards of Agricultural and Forest Policy, Federal Ministry of Agriculture and Nutrition (2016) Klimaschutz in der Land- und Forstwirtschaft sowie den nachgelagerten Bereichen Ernährung und Holzverwendung. Berlin, Germany (in German)

Slot M, Cala D, Aranda J, Virgo A, Michaletz ST, Winter K (2021) Leaf heat tolerance of 147 tropical forest species varies with elevation and leaf functional traits, but not with phylogeny. Plant Cell Environ 44:2414–2427

Spiecker H, Lindner M, Schuler J (eds) Douglas-Fir - an option for Europe What Science Can Tell Us vol. 9. European Forest Institute, Joensuu

Still CJ, Page G, Rastogi B et al. (2022) No evidence of canopy-scale leaf thermoregulation to cool leaves below air temperature across a range of forest ecosystems. Proc Natl Acad Sci USA 119:e2205682119

Teskey R, Wertin T, Bauweraerts I, Ameye M, McGuire MA, Steppe K (2015) Tree response to extreme heat. Plant Cell Environ 38:1699–1712

Thomas FM, Rzepecki A, Werner W (2022) Non-native Douglas fir (*Pseudotsuga menziesii*) in Central Europe: ecology, performance and nature conservation. For Ecol Manag 506:119956

Tiwari R, Gloor E, da Cruz WJA et al. (2021) Photosynthetic quantum efficiency in south-eastern Amazonian trees may be already affected by climate change. Plant Cell Environ 44:2428–2439

van Tiel N, Lenczner G, Rao MP, Grossiord C, Tuia D (2026) Tropical forests are facing increasing risks of exposure to critical temperature thresholds. Proc Natl Acad Sci USA 123:e2528622123

Visakorpi K, Manzanedo RD, Görlich AS, Schiendorfer K, Altermatt Bieger A, Gates E, Hille Ris Lambers J (2024) Leaf-level resistance to frost, drought and heat covaries across European temperate tree seedlings. J Ecol 112:559–574

Winter K, Krüger Nuñez CR, Slot M, Virgo A (2025) In thermotolerance tests of tropical tree leaves, the chlorophyll fluorescence parameter *F*_v_/*F*_m_ measured soon after heat exposure is not a reliable predictor of tissue necrosis. Plant Biol 27:146–153

WMO (2026) Western Europe has hottest June on record. World Meteorological Organization, Geneva, Switzerland. https://wmo.int/media/news/western-europe-has-hottest-june-record (accessed on 22 July 2026)

Zhang H, Ning Q, Li Q, Jin Y, Cao Y, Bakpa EP, Zhao H, Song J, Ye P, Wen Y, Song L, Liu, H (2024) Contrasting heat tolerance of evergreen and deciduous urban woody species during heat waves. Funct Ecol 38:1649–1660

